# All-optical electrophysiology refines populations of in silico human iPS-CMs for drug evaluation

**DOI:** 10.1101/799478

**Authors:** M Paci, E Passini, A Klimas, S Severi, J Hyttinen, B Rodriguez, E Entcheva

## Abstract

High-throughput *in vitro* drug assays have been impacted by recent advances in human induced pluripotent stem cell-derived cardiomyocytes (hiPS-CMs) technology and by contact-free all-optical systems simultaneously measuring action potential (AP) and Ca^2+^ transient (CaTr). Parallel computational advances have shown that *in silico* models can predict drug effects with high accuracy. In this work, we combine these *in vitro* and *in silico* technologies and demonstrate the utility of high-throughput experimental data to refine *in silico* hiPS-CM populations, and to predict and explain drug action mechanisms. Optically-obtained hiPS-CM AP and CaTr were used from spontaneous activity and under pacing in control and drug conditions at multiple doses.

An updated version of the Paci2018 model was developed to refine the description of hiPS-CM spontaneous electrical activity; a population of *in silico* hiPS-CMs was constructed and calibrated using the optically-recorded AP and CaTr. We tested five drugs (astemizole, dofetilide, ibutilide, bepridil and diltiazem), and compared simulations against *in vitro* optical recordings.

Our simulations showed that physiologically-accurate population of models can be obtained by integrating AP and CaTr control records. Thus constructed population of models predicted correctly the drug effects and occurrence of adverse episodes, even though the population was optimized only based on control data and *in vitro* drug testing data were not deployed during its calibration. Furthermore, the *in silico* investigation yielded mechanistic insights, e.g. through simulations, bepridil’s more pro-arrhythmic action in adult cardiomyocytes compared to hiPS-CMs could be traced to the different expression of ion currents in the two.

Therefore, our work: i) supports the utility of all-optical electrophysiology in providing high-content data to refine experimentally-calibrated populations of *in silico* hiPS-CMs, ii) offers insights into certain limitations when translating results obtained in hiPS-CMs to humans and shows the strength of combining high-throughput *in vitro* and population *in silico* approaches.

**Significance:** We demonstrate the integration of human *in silico* drug trials and optically-recorded simultaneous action potential and calcium transient data from human induced pluripotent stem cell-derived cardiomyocytes (hiPS-CMs) for prediction and mechanistic investigations of drug action. We propose a population of *in silico* models i) based on a new hiPS-CM model recapitulating the mechanisms underlying hiPS-CM automaticity and ii) calibrated with all-optical measurements. We used our *in silico* population to predict and evaluate the effects of 5 drugs and the underlying biophysical mechanisms, obtaining results in agreement with our experiments and one independent dataset. This work supports the use of high-content, high-quality all-optical electrophysiology data to develop, calibrate and validate computer models of hiPS-CM for *in silico* drug trials.

## Introduction

Both, new *in silico* methods and the use of human induced pluripotent stem cell-derived cardiomyocytes (hiPS-CMs) have become increasingly important in tackling the challenge of assessment and prediction of drug effects and their potential cardiotoxicity, as supported by the Comprehensive In Vitro Proarrhythmia Assay (CiPA) initiative (1, 2). Many *in silico* studies on this topic have been published in recent years, showcasing a variety of methodologies, including electrophysiological models of cardiac cells, machine learning algorithms, and a combination of both (3–8). The potential of hiPS-CMs for drug-induced pro-arrhythmia predictions *in vitro* has been shown in many experimental studies (9, 10) despite certain outstanding limitations. Concerns lie with their high inter-lab and inter-batch variability and level of maturity compared to adult cardiomyocytes (11), e.g. spontaneous beating, cell morphology, disorganization of their contractile elements (12), and different ion channel expression (13). Nevertheless, hiPS-CMs represent the best experimental platform to date to study human cardiac electrophysiology and drug action in a rigorous and scalable/high-throughput way. *In silico* models of hiPS-CMs have emerged (14–17) as an invaluable tool to better understand the distinct ionic mechanisms underlying hiPS-CM’s drug response (18, 19). The robustness of *in silico* models depends on the amount and the quality of the experimental data used in their calibration and validation. Traditionally, such data have been acquired from a limited number of isolated cells (outside of their multicellular environment), through time-demanding and tedious manual patch-clamp techniques.

Limited experimental data present challenges of not being able to capture the genotypical and the phenotypical variability observed in a cell population, which is especially relevant for the highly-variable hiPS-CMs. These challenges have been partially addressed through modelling and data curation. *In silico* population of models approaches have been developed to reflect the wider range of parameters beyond the limited experimental data (20, 21). Database merging has also been used in the desire to expand the experimental data needed to tune the model parameters, e.g. in (19, 22) we merged 6 *in vitro* datasets of action potential (AP) biomarkers to generate a population of *in silico* hiPS-CMs. Using data from different laboratories widen the data variability considerably.

On the technology side, the problem of limited experimental data has been tackled by new experimental techniques with increased throughput and amenable to automation, e.g. automated patch-clamp platforms (23, 24) or microelectrode arrays (MEAs) (13). However, these techniques still suffer the limitations of probe-sample physical contact, which limits their performance with hiPS-CMs (25). Contact-free optical recordings overcome these limitations and offer comprehensive characterization. Calcium and contraction-measurement systems have been leveraged for cardiotoxicity testing (26). Ahola et al. (27, 28) developed a video-based contact-free method to quantify the biomechanics of beating hiPS-CMs, by processing simultaneous recording of motion and Ca^2+^ transients (CaTr) from fluorescence videos. However, AP signals represent key aspects of cardiotoxicity responses that may not be captured by field potentials, CaTr or mechanical contractions. All-optical electrophysiology (29, 30) approaches offer contactless interrogation and high-throughput records of voltage and calcium in an attempt to increase information content. Application of these techniques to drug screening with hiPS-CMs have been successfully demonstrated (25, 31, 32), including our OptoDyCE that combines optical pacing and simultaneous optical records of voltage and calcium or contractions. The use of optical systems with hiPS-CMs preparations provides an abundance of *in vitro* data with the potential to provide an excellent basis to construct experimentally-calibrated population of *in silico* hiPS-CMs. The value of such high-content optical recordings of CaTr and AP (without ion channel level data) to constrain *in silico* populations of models remains to be tested.

The main goal of this work was to demonstrate the utility of *in silico* simulation trials informed by all-optical cardiac electrophysiology (optically-obtained high throughput measurements of AP and CaTr from hiPS-CMs under spontaneous and optically-triggered conditions) for prediction and mechanistic understanding of drug action. Optically-obtained AP and CaTr measurements are used to guide and improve the design and calibration of a population of *in silico* hiPS-CMs. We then test the performance of *in silico* simulation trials with the populations of models against *in vitro* drug trials for 5 reference compounds, both in terms of their consistency and to deepen the mechanistic insights unravelled. In detail: i) We present an improved version of the Paci2018 hiPS-CM model (15), providing improved simulation of the Na^+^/Ca^2+^ exchanger (I_NCX_) role in sustaining the automaticity of AP. ii) We use high throughput optical measurement of AP, CaTr alone and both to calibrate an *in silico* population of hiPS-CMs models. iii) We challenge this population by applying 5 reference compounds at multiple concentrations, and comparing the results against *in vitro* data, not used for the calibration step. We investigate the mechanisms underlying the different response to bepridil in hiPS-CMs (both *in vitro* and *in silico*) compared to adult cardiomyocytes.

## Materials and Methods

### Experimental dataset

The experimental dataset consists of AP and CaTr recordings from hiPS-CMs syncytia (CDI iCell^2^ cardiomyocytes) obtained with the all-optical OptoDyCE system (25) in a 384-well plate format at room temperature (21°C) and with extracellular concentrations Nao = 135.0, Ko = and Cao = 1.33mM, in both paced and non-paced conditions. Recordings were performed in control conditions (0.1% DMSO) and following application of 5 reference compounds: astemizole (antihistamine), dofetilide (antiarrhythmic agent, class III), ibutilide (antiarrhythmic agent, class III) bepridil (antiarrhythmic agent, class IV) and diltiazem (antiarrhythmic agent, class IV).

Control recordings were performed on 10 plates (384-well format). Voltage and calcium-derived biomarkers were obtained from 5 independent multicellular samples per plate (each sample having at least 200 cells). The following biomarkers were considered: AP and CaTr cycle length (AP CL and CaTr CL), duration at 30%, 50% and 90% of AP repolarization (APD_30_, APD_50_ and APD_90_) and of CaTr decay (CTD_30_, CTD_50_, CTD_90_), AP and CaTr triangulation (AP Tri_90-30_=APD_90_-APD_30_ and CaTr Tri_90-30_=CTD_90_-CTD_30_) and CaTr time from CaTr onset to peak (CaTr tRise_0,peak_). Each measurement was characterized by its mean value (mean) and its standard deviation (SD) over a variable number of beats for each multicellular sample. Some acquisitions failed, and were discarded from the dataset, leading to a total of 42 control non-paced and 49 control paced multicellular samples, thus integrating responses from over 8400 cells. Min and Max experimental ranges for each biomarker were computed by defining a lower and upper bounds (*LB* = *min*(*mean* − 2 ∗ *SD*) and *UB* = *max*(*mean* − 2 ∗ *SD*), respectively), for non-paced and paced measurements, as reported in Table 1.

**Table 1.** Experimental ranges of the *in vitro* optical recordings.

|  | Control non-paced |  | Control paced |  |
| --- | --- | --- | --- | --- |
|  | Lower Bound | Upper Bound | Lower Bound | Upper Bound |
|  | (LB<sub>NP</sub>) | (UB<sub>NP</sub>) | (LB<sub>P</sub>) | (UB<sub>P</sub>) |
| AP CL (ms) | 1310.2 | 12,798.5 | --- | --- |
| ADP<sub>90</sub> (ms) | 485.1 | 1393.8 | 514.3 | 1397.6 |
| APD<sub>50</sub> (ms) | 310.6 | 1059.6 | 332.5 | 932.4 |
| APD<sub>30</sub> (ms) | 240.6 | 910.3 | 261.6 | 786.3 |
| AP Tri<sub>90-30</sub> (ms) | 132.0 | 741.0 | 251.9 | 839.9 |
| CaTr CL (ms) | 1310.1 | 12,805.3 | --- | --- |
| CTD<sub>90</sub> (ms) | 754.9 | 2897.5 | 863.2 | 1803.0 |
| CTD<sub>50</sub> (ms) | 463.8 | 1376.4 | 510.5 | 1167.8 |
| CTD<sub>30</sub> (ms) | 353.4 | 1065.2 | 382.7 | 983.7 |
| CaTr tRise<sub>0,peak</sub> (ms) | 112.9 | 622.3 | 96.5 | 473.8 |
| CaTr Tri<sub>90-30</sub> (ms) | 104.8 | 1872.5 | 438.6 | 1140.6 |

LB<sub>NP</sub>: lower bound, non-paced; UB<sub>NP</sub>: upper bound, non-paced; LB<sub>P</sub>: lower bound, paced; UB<sub>P</sub>: upper bound, paced; see main text for biomarker descriptions.

Reference compounds were tested in 5 plates (one for each drug), considering 4 increasing doses (D1, D2, D3 and D4) and 6 multicellular samples for each dose (thus integrating responses from at least 1200 cells per drug dose). After discarding failed recordings, we used the same methods as in control to compute the experimental biomarker ranges.

### Updated version of the Paci2018 hiPS-CM model

A limitation of the Paci2018 hiPS-CM model (15) was noted - namely, failure to reproduce the cessation of the spontaneous electrical activity following strong block of the I_NCX_, as shown by recent *in vitro* and *in silico* experiments (16, 33). A very large window current in Paci2018 for the fast Na^+^ current (I_Na_) was identified as the key to sustaining the automaticity upon I_NCX_ block. We improved the Paci2018 model to reproduce this specific mechanism, while preserving all its good features. We kept the same structure of the Paci2018: the model includes two compartments, namely cytosol and sarcoplasmic reticulum (SR), and it follows the classical Hodgkin & Huxley formulation, which describe the membrane potential as

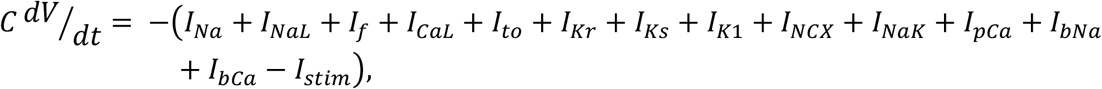

where *C* is the membrane capacitance *V* the membrane voltage and *I*_*stim*_ the stimulus current. The ion current/pumps in the model are: I_Na_, the late Na^+^ current (I_NaL_), the funny current (I_f_) the L-type Ca^2+^ current (I_CaL_), the transient outward K^+^ current (I_to_), the rapid and slow delayed rectifier K^+^ currents (I_Kr_ and I_Ks_), the inward rectifier K^+^ current (I_K1_), the Na^+^/Ca^2+^ exchanged (I_NCX_), the Na^+^/K^+^ pump (I_NaK_) the sarcolemmal Ca^2+^ pump (I_pCa_) and the Na^+^ and Ca^2+^ background currents (I_bNa_ and I_bCa_). The SR compartment exchanges Ca^2+^ with cytosol through three fluxes: RyR-sensitive release current (I_rel_), SERCA pump (I_up_) and leakage current (I_leak_).

To develop the Paci2019 model (details in the Supporting Material):

- we updated the formulations for I_Na_ and I_f_ with the ones proposed in (16);
- we optimized the model parameters to fit the same dataset of *in vitro* AP and CaTr biomarkers used for (15), which have been recorded at 37°C;
- we validated the model against the same experimental protocols used for (15).

As a result, we obtained an improved version of our hiPS-CM model (Paci2019), where the spontaneous electrical activity is triggered both by I_f_ and Ca^2+^ release from the sarcoplasmic reticulum, which in turn depolarize the membrane potential via I_NCX_. Details on the optimization procedure are reported in the Supporting Material, together with the model parameter values and equations.

The optically-obtained *in vitro* data in this paper were recorded at 21°C and with extracellular concentrations (Nao = 135.0, Ko = 5.4 and Ca_o_ = 1.33mM instead of Nao = 150.0, Ko = 5.4 and Cao = 1.8mM). Consequently, we implemented temperature correction of the new Paci2019 model to these conditions. Temperature difference was managed by setting the correct temperature in the specific model parameter affecting the Nernst potentials and ion currents such as I_NCX_ or I_NaK_, rescaling the time constants of the other main ionic currents by means of the Q_10_ factors reported in (34–37), and summarized in Table S1 in the Supporting Material.

### hiPS-CM *in silico* population calibrated with optical AP and CaTr recordings

The new Paci2019 model, adapted to the temperature and extracellular concentrations of the optical recordings, was used as baseline to construct a population of *in silico* hiPS-CMs, based on the population of models methodology (19, 20, 38). We sampled a total of 22 parameters in the [50-150]% range compared to their original values. Parameters were chosen similarly to (39), to include all the main ionic conductances, as well as key kinetics parameter, known to impact both AP and CaTr biomarkers: (i) the maximum conductances of I_Na_, I_NaL_, I_f_, I_CaL_, I_to_, I_Ks_, I_Kr_, I_K1_, I_NCX_, I_NaK_, I_pCa_, I_rel_, I_up_; (ii) activation and inactivation time constants of I_Na_, I_CaL_ and I_rel_; (iii) adaptation time constant and half inactivation Ca^2+^ concentration of I_rel_; (iv) I_up_ half saturation constant. An initial population of 30,000 hiPS-CMs was generated, and then calibrated based on the optical recordings, i.e. only the models whose biomarkers were in agreement with the *in vitro* data were maintained. Biomarkers were computed in steady state (after 800s), as the average on the last 20 beats. The lack of absolute amplitude values for AP in the optically-recorded data was handled by an additional biomarker to constrain the amplitude of the non-paced AP (AP peak between 17.0 and 57.7 mV), as in (19).

Three different calibration options were performed considering both paced and non-paced biomarkers, thus generating three different experimentally-calibrated populations: i) All AP and CaTr biomarkers (AP_CaTr population); ii) AP biomarkers only (AP_only population); iii) CaTr biomarkers only (CaTr_only population). The three populations were compared to investigate how the choice of AP and CaTr biomarkers affect the calibration process and the coverage of the biomarker space compared to experimental ranges.

### In silico drug trials

In silico drug trials were performed for 5 compounds (astemizole, dofetilide, ibutilide, bepridil and diltiazem) considering the 4 concentrations for each tested *in vitro*. Drug simulations were run for 400s from steady state conditions. Models were not paced, to also investigate drug-induced effect on the spontaneous beating frequency. We used a simple pore-block drug model as in (3, 19, 38), consisting of IC_50_ and Hill’s coefficients from literature and reported in Table S2 in the Supporting Material. The experimental concentrations for each drug are reported in Table S3 in the Supporting Material, together with the corresponding percentage of residual currents following drug application and the maximal effective free therapeutic concentration (EFTPC_max_), for comparison.

Because of the discrepancy between hiPSC and adult CMs observed for bepridil ((40) vs (3, 41)), only for bepridil 10μM, we run additional tests, reducing its I_CaL_ blocking action to half (64% residual I_CaL_ instead of 32%) and to zero (100% residual I_CaL_), while preserving its blocking action on the other ion channels. This test was done on 4 models selected among the ones that showed a pro-arrhythmic behaviour when administered astemizole.

Following drug application, we assessed the drug-induced changes on AP and CaTr biomarkers, as well as the occurrence of abnormalities. Single and multiple early after-depolarizations (EADs) were defined as extra-peaks greater than −55mV in between two consecutive AP upstrokes. Repolarisation failure were identified when a stable (dV/dt_max_<0.1 V/s) membrane potential > −40 mV was observed during the last 15 s of simulation. Irregular rhythm was identified when the difference in cycle length between two consecutive AP greater than 150%.

We looked also for two additional phenotypes, that we did not consider as abnormalities: quiescence (40) and residual activity (42), mainly occurring during diltiazem administration (see Results). If a model reacted to drug by producing AP whose peaks were greater than −40mV but smaller than 0mV, we labelled the model as residual activity. Conversely, we considered the model quiescent, i.e. not producing anymore spontaneous AP, if during the last 15s the average membrane potential was smaller than −40mV or a potential residual activity had all the peaks smaller than −40mV.

## Results

### The new Paci2019 hiPS-CMs model

The automated optimization process successfully identified a new Paci2019 model in agreement with the *in vitro* AP and CaTr biomarkers used in (15), as shown in Table 2. Figure S1 in the Supporting Material shows a detailed comparison between the new model (in black) and the Paci2018 model (in red) (15). Parameter values are reported in the Supporting Material.

**Table 2.** Action potential and Ca^2+^ transient biomarkers simulated by the Paci2019 hiPS-CM model at 37°C

| Biomarker<br>(reference) | Experimental value<br>(mean±SD) | Simulated<br>value |
| --- | --- | --- |
| APA (mV) (43) | 104 ± 6.0 | 102.0 |
| MDP (mV) (43) | -75.6 ± 6.6 | -74.9 |
| AP CL (ms) (43) | 1700.0 ± 547.7 | 1712.4 |
| dV/dt<sub>max</sub> (V/s) (43) | 27.8 ± 26.3 | 20.4 |
| APD<sub>10</sub> (ms) (43) | 74.1 ± 26.3 | 87.0 |
| APD<sub>30</sub> (ms) (43) | 180 ± 58.6 | 223.8 |
| APD<sub>90</sub> (ms) (43) | 414.7 ± 119.4 | 390.2 |
| AP Tri (-)(43) | 2.5 ± 1.1 | 2.8 |
| CaTr DURATION (ms) (15) | 804.5 ± 188.0 | 691.5 |
| CaTr tRise<sub>10,50</sub> (ms) (15) | 82.9 ± 50.5 | 55.0 |
| CaTr tRise<sub>10,90</sub> (ms) (15) | 167.3 ± 69.8 | 118.2 |
| CaTr tRise<sub>10,peak</sub> (ms) (15) | 270.4 ± 108.3 | 184.0 |
| CaTr tDecay<sub>90,10</sub> (ms) (15) | 409.8 ± 100.1 | 341.0 |
| CaTr CL (ms) (15) | 1653.9 ± 630 | 1712.4 |

AP and CaTr biomarkers from (15). Both AP and CaTr biomarkers were recorded at 37°C. AP biomarkers (patch-clamp): AP amplitude (APA); maximum diastolic potential (MDP); cycle length (CL); maximum upstroke velocity (dV/dt<sub>max</sub>); AP duration at 10%, 30% and 90% of repolarization (APD<sub>10</sub>, APD<sub>30</sub>, APD<sub>90</sub>); AP triangulation (AP Tri) computed as the ratio between APD<sub>30</sub>-APD<sub>40</sub> and APD<sub>70</sub>-APD<sub>80</sub>; CaTr DURATION; CaTr rise time from 10% to 50% (CaTr tRise<sub>10,50</sub>), 90% (CaTr tRise<sub>10,90</sub>), and to CaTr peak (CaTr tRise<sub>10,peak</sub>), decay time from 90% to 10% (CaTr tDecay<sub>90,10</sub>), and CaTr rate (CaTr CL).

The main difference between the two models is the shape of the I_NCX_ current. Before the upstroke, the new I_NCX_ provides an additional inward contribution (−0.5A/F) that is added to If (−0.25A/F), supporting the membrane depolarization and allowing the opening of the I_Na_ channels. Figure 1 illustrates the contribution of I_NCX_ to the hiPS-CM automaticity, as reported in (16, 33): blocking I_NCX_ reduces its inward component slowing down the rate of spontaneous AP, up to suppression. In particular, an issue in the Paci2018 model was that AP suppression did not happen, in disagreement with *in vitro* data by Kim et al. (33) in response to 2μM SEA0400, an inhibitor of the forward I_NCX_ in a cluster of hiPS-CMs. The large I_Na_ window current was identified as a key factor in supporting the automaticity, thus making the Paci2018 model unable to capture the aforementioned mechanism.

**Figure 1.**
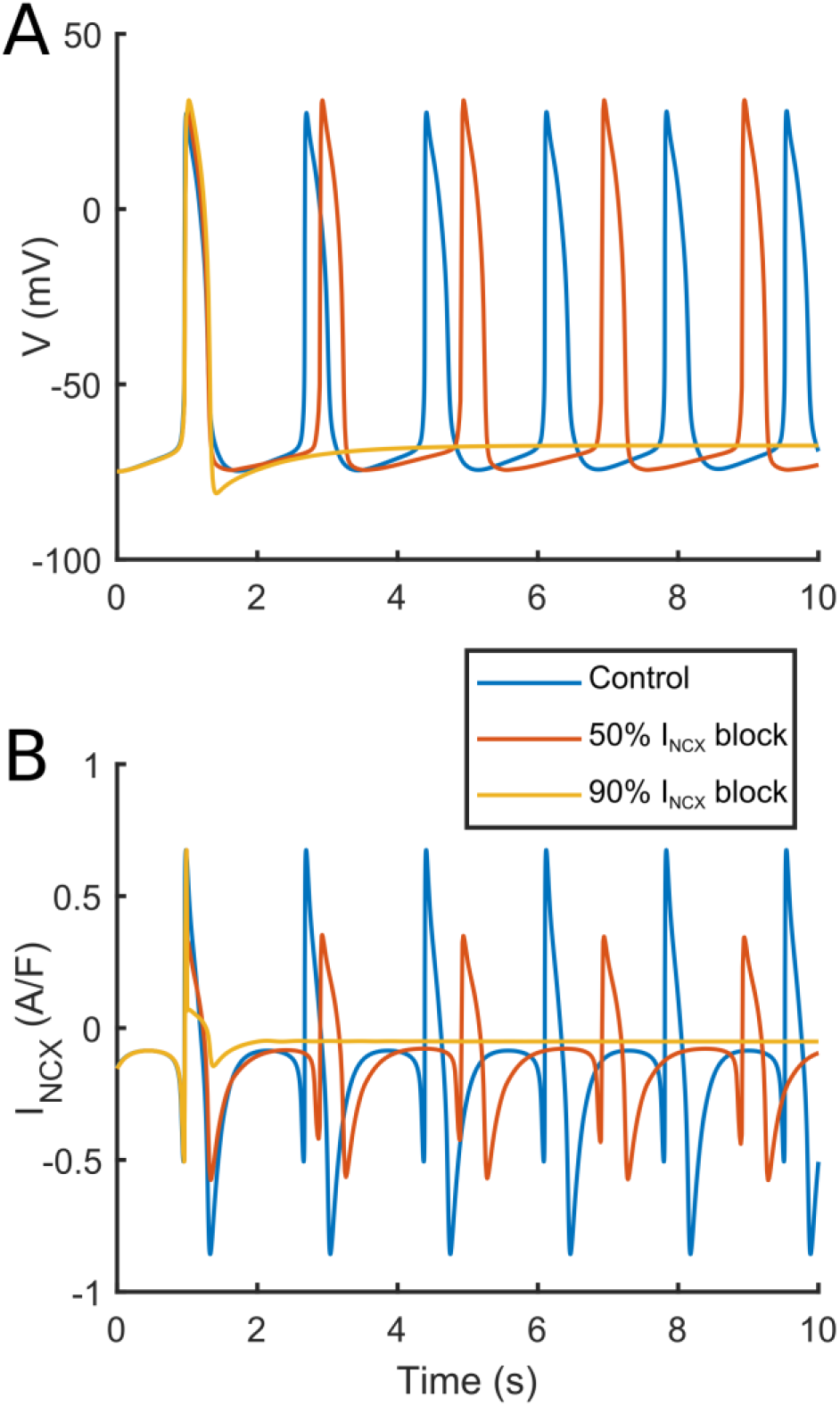
Effects of different levels of I_NCX_ block on the spontaneous AP simulated using the Paci2019 in control (blue line), with 50% I_NCX_ block (red line), and suppressed when considering high I_NCX_ block (yellow line).

The new Paci2019 model can simulate spontaneous Ca^2+^ release from the SR both with standard extracellular Ca^2+^ concentration (Ca_o_ = 1.8mM, Figure S2 in the Supporting Material) and Ca^2+^ overload (simulated by increasing the extracellular Ca^2+^ concentration to Ca_o_ = 2.8, 2.9 and 3.0mM, Figure S3 in the Supporting Material). Moreover, it reproduces well the *in vitro* data by Ma et al. (43) with ion channel blockers (Figure S4 in the Supporting Material), I_f_ block and hyperkalemia experiments as (33) (see Supporting Information) and alternans in ischemia-like conditions as (15) (Figure S5 in the Supporting Material). Finally, the CaTr amplitude of 160 nM is in agreement with data by Rast et al. (44), recorded from hiPS-CM ensembles incubated at 37°C (calibrated Fura-2-based photometry measures) and not used for model calibration.

After updates for the extracellular ion concentrations and temperature adjustment, as described in Methods, the Paci2019 model’s AP and CaTr biomarkers moved closer to the optical recordings reported in Table 1 obtained at room temperature. For example, spontaneous CL increased (from 1,712 to 4,144 ms) and APD_90_ prolonged from 390 to 1,119 ms. Figure 2 shows a comparison of the Paci2019 model (green traces) vs. the same model adapted for extracellular concentrations and temperature (blue traces).

**Figure 2.**
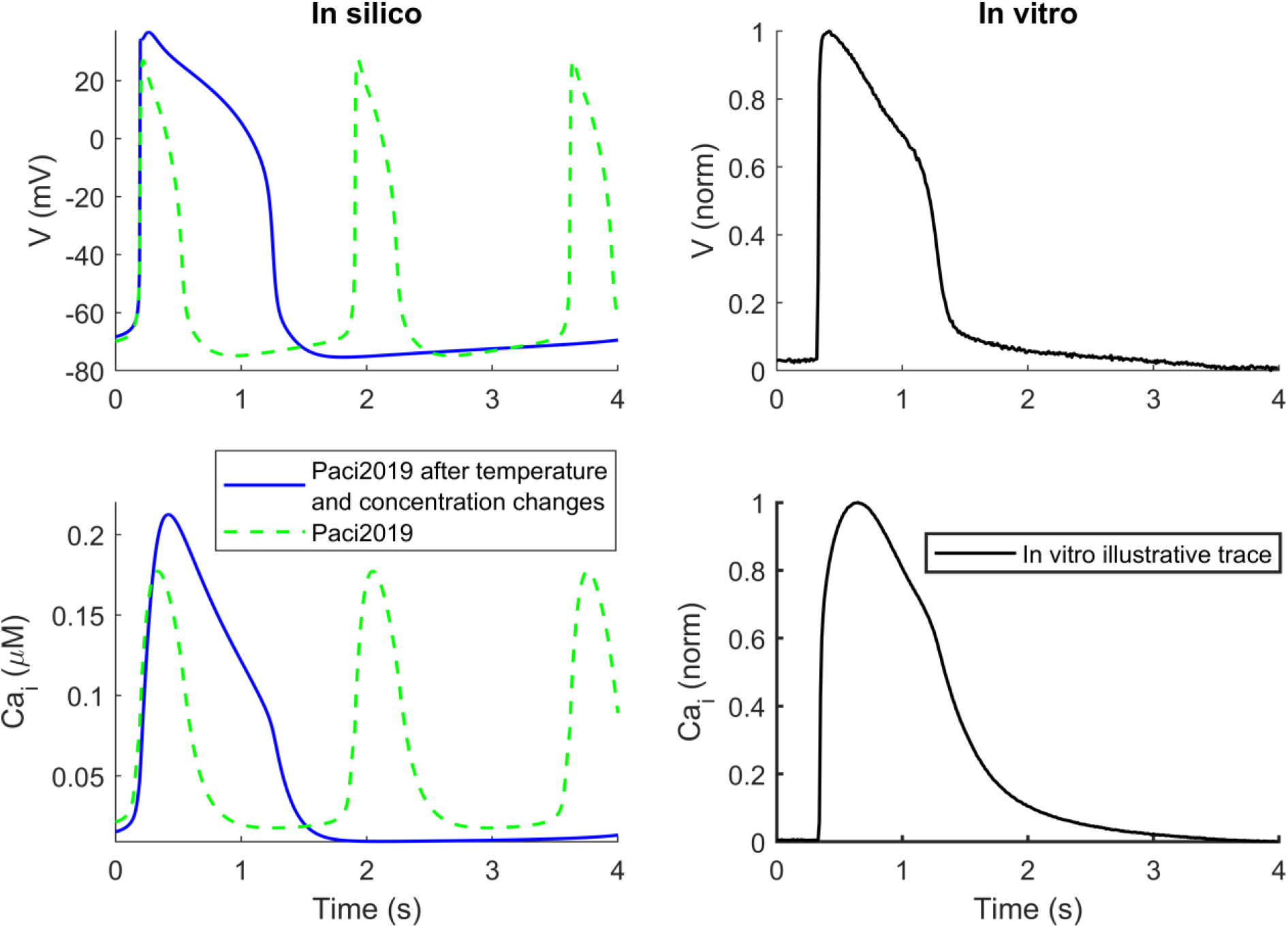
Simulated spontaneous AP and CaTr for the Paci2019 model at 37°C (green) vs the same model adapted to 21°C (blue) and extracellular concentrations as in the *in vitro* optical recordings (right column, spontaneous illustrative trace).

### Single dataset calibration vs combined dataset calibration

The Paci2019 model, adapted for the extracellular concentrations and room temperature used in the *in vitro* experiments, was deployed to generate an initial population of 30,000 models. As described in Methods, 3 different calibrations were performed (using AP only, CaTr only or both AP and CaTr biomarkers), leading to 3 calibrated populations: AP_only, CaTr_only and AP_CaTr, respectively.

A comparison of the AP and CaTr biomarkers for the 3 populations is shown in Figure 3. The AP_only population (green boxplots) consists of 969 models. As expected, it shows good agreement with the experimental AP biomarkers in addition to a good coverage of the experimental ranges, both non-paced and paced (Panel A and B). However, many models have CaTr biomarkers outside the experimental ranges, e.g. CTD_90_, CTD_50_ and CTD_30_ are often too short (Panel C and D). The CaTr_only population (black boxplots) consists of 5,030 models in good agreement with CaTr biomarkers, both non-paced and paced (Panels C and D). However, many models yield AP durations and triangulation outside the experimental ranges (Panels A and B). As expected, the AP_CaTr population, obtained by calibrating with both AP and CaTr biomarkers (blue boxplots), appears to be the best constrained, with 477 models showing good agreement and coverage of the biomarker space.

**Figure 3.**
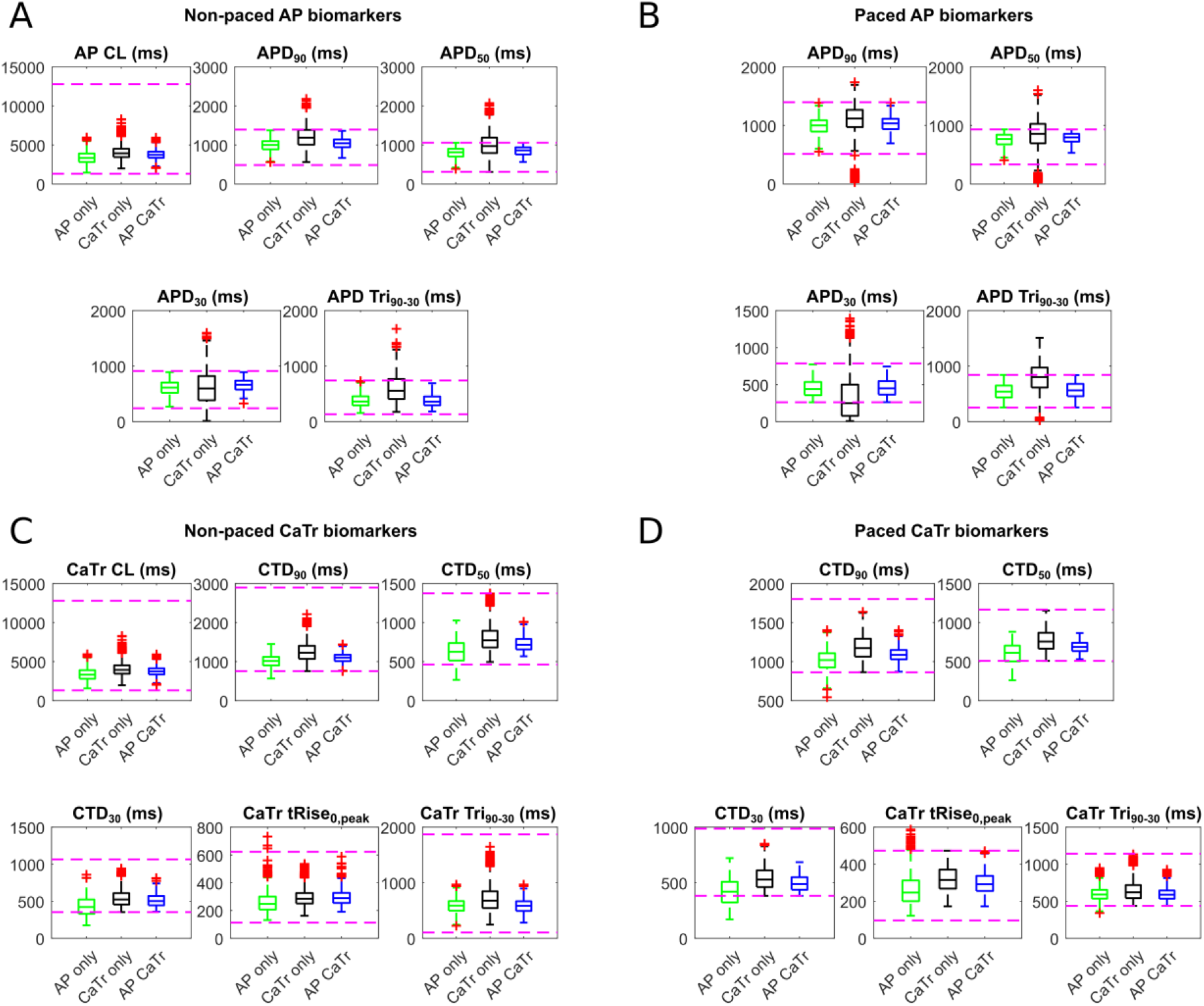
AP (A, B) and CaTr (C, D) biomarker distributions in the three populations of hiPS-CM models, calibrated with *in vitro* AP biomarkers only (green), CaTr biomarkers only (black) or both (blue). On each box, the central mark is the median of the population, box limits are the 25^th^ and 75^th^ percentiles, and whiskers extend to the most extreme data points not considered outliers. Red crosses represent outliers. The dashed magenta lines represent the lower and upper bounds of the experimental recordings, as reported in Table 1.

Figure 4 shows the distributions of the seven parameters with differential responses in the 3 experimentally-calibrated populations (|Δ*median*| > 10% between AP_only/CaTr_only and AP_CaTr). Distributions of all parameters varied in the population are shown in Figure S6 in the Supporting Material. Adding AP biomarkers for calibration (AP_only and AP_CaTr populations vs. CaTr_only) helps adjust five key parameters in important ways (lowers their median values): G_Na_, I_Na_ inactivation time constants, G_K1_, I_NCX_ maximum current and I_CaL_ inactivation time constant (Figure 4). The smaller G_Na_ is due to the upper limit on AP peak. This also imposes a smaller I_Na_ inactivation time constant (faster inactivation), further contributing to reduced AP peak amplitude. A lower G_K1_ results in a slightly depolarized MDP, consequently reducing I_Na_ availability, and again limiting the AP peak. A reduced I_NCX_ maximum current prevents an excessively fast early repolarization phase, e.g. short APD_30_. Finally, a smaller I_CaL_ inactivation time constant speeds up I_CaL_ inactivation, thus limiting excessively long AP.

**Figure 4.**
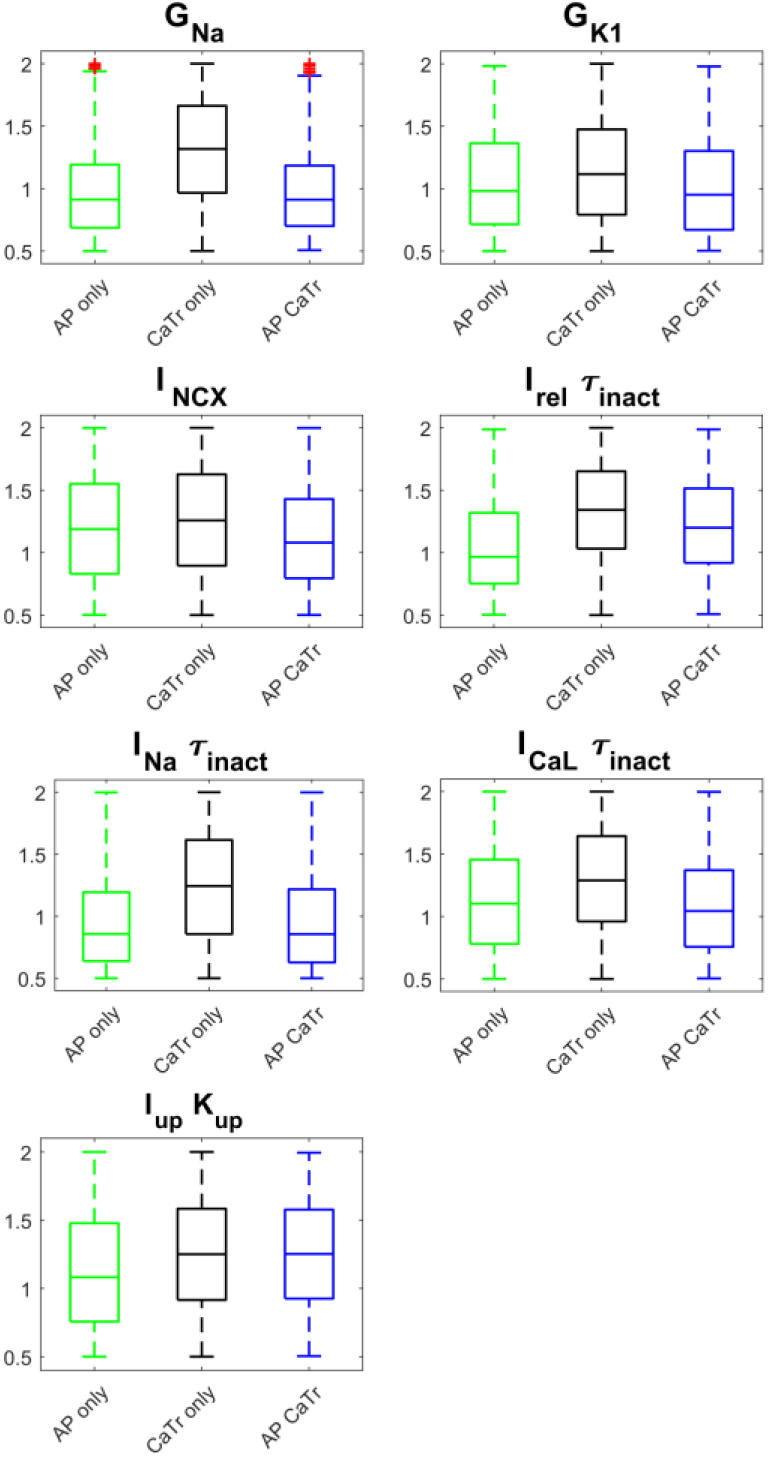
Parameter distributions for the three populations: AP_only (green), CaTr_only (black) and AP_CaTr (blue). Red crosses represent outliers. Boxplot description as in Figure 3. Only parameters with |Δmedian|>10% between AP_only/CaTr_only and AP_CaTr are shown here, while distributions of all 22 parameters are reported in Figure S6.

Considering CaTr biomarkers for calibration (CaTr_only and AP_CaTr, vs AP_only) increases the median values for two calcium-release parameters: I_rel_ inactivation time constant and I_up_ half saturation value (Figure 4). The first causes a slower inactivation of I_rel_, and consequently a longer CaTr (Figure 3, Panels C-D). The latter, that appears in the denominator of the I_up_ formulation (15), causes a reduction of Ca^2+^ uptake, thus also contributing to a longer CaTr.

Overall, these results reveal important information contributed by the AP or CaTr biomarkers in the calibration process to better capture the experimental recordings. For the rest of this study, including the *in silico* drug trials, only the AP_CaTr population of 477 hiPS-CM models was considered. The AP and CaTr traces for this population are shown in Figure 5.

**Figure 5.**
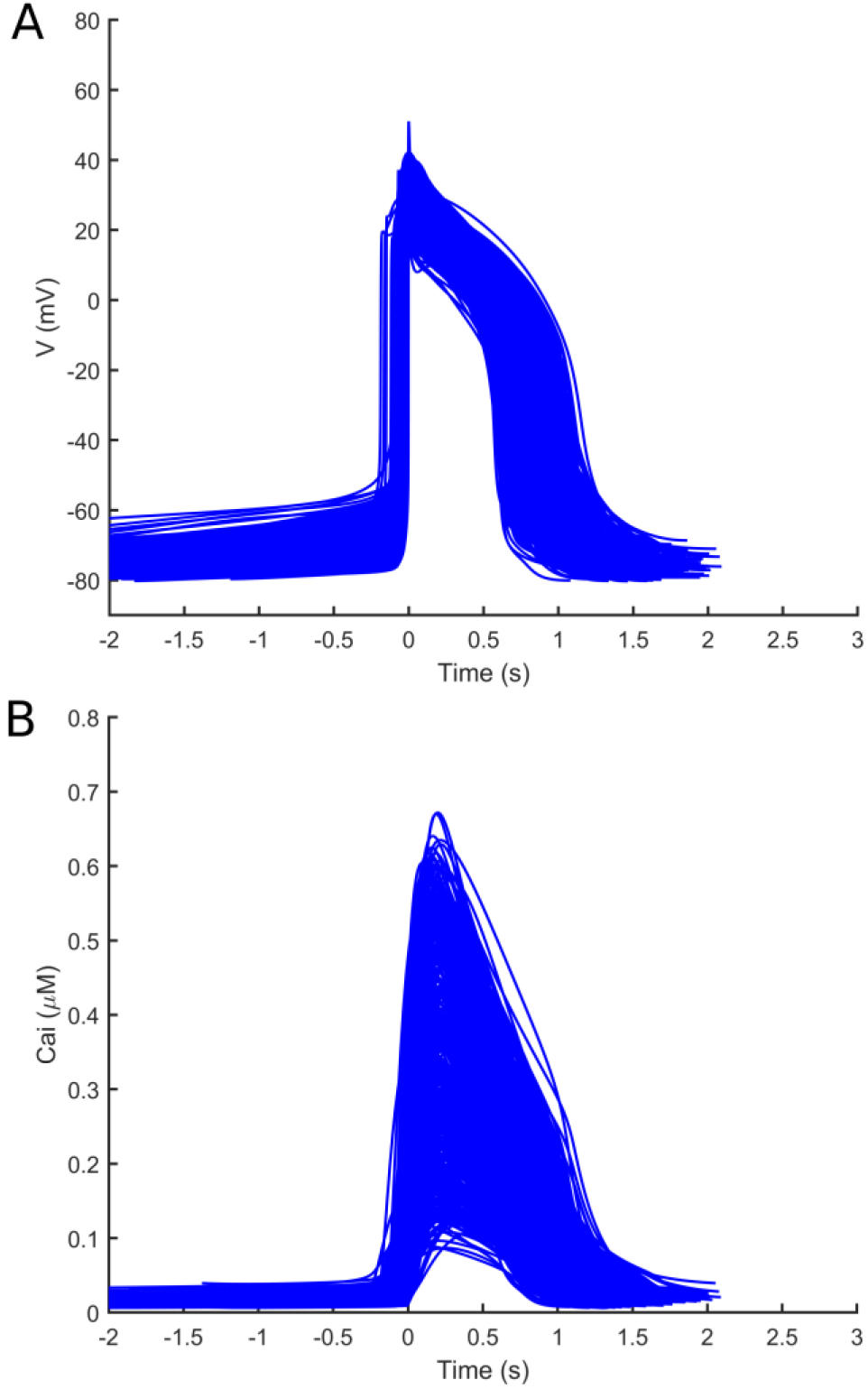
AP (A) and CaTr (B) traces included in the final population of 477 *in silico* hiPS-CMs, calibrated with both AP and CaTr biomarkers.

### In silico drug trials

Using the population of 477 hiPS-CM models shown in Figure 5, calibrated with both experimental AP and CaTr biomarkers, we ran *in silico* drug trials for 5 reference compounds (astemizole, dofetilide, ibutilide, bepridil, diltiazem) at 4 increasing concentrations (D1-D4) each. Simulation results were validated against the corresponding *in vitro* experiments, which were not used during the calibration process. For each drug trial, we checked how the drug affects the AP and CaTr biomarkers compared to control (D0), and assessed the presence of drug-induced abnormalities. Figure 6 summarizes the drug effects on four AP and CaTr biomarkers (AP CL, APD_90_, CTD_90_ and CaTr Tri_90-30_). Shown are: i) *in silico* biomarker boxplots for the models that after drug administration still produce spontaneous AP and CaTr at room temperature and at the ion concentrations tested *in vitro*; and ii) *in vitro* optically-recorded biomarkers (green/purple diamonds) and their variability ranges (green/purple bars). Results for all biomarkers are shown in the Supporting Information, Figure S7-11 in the Supporting Material.

**Figure 6.**
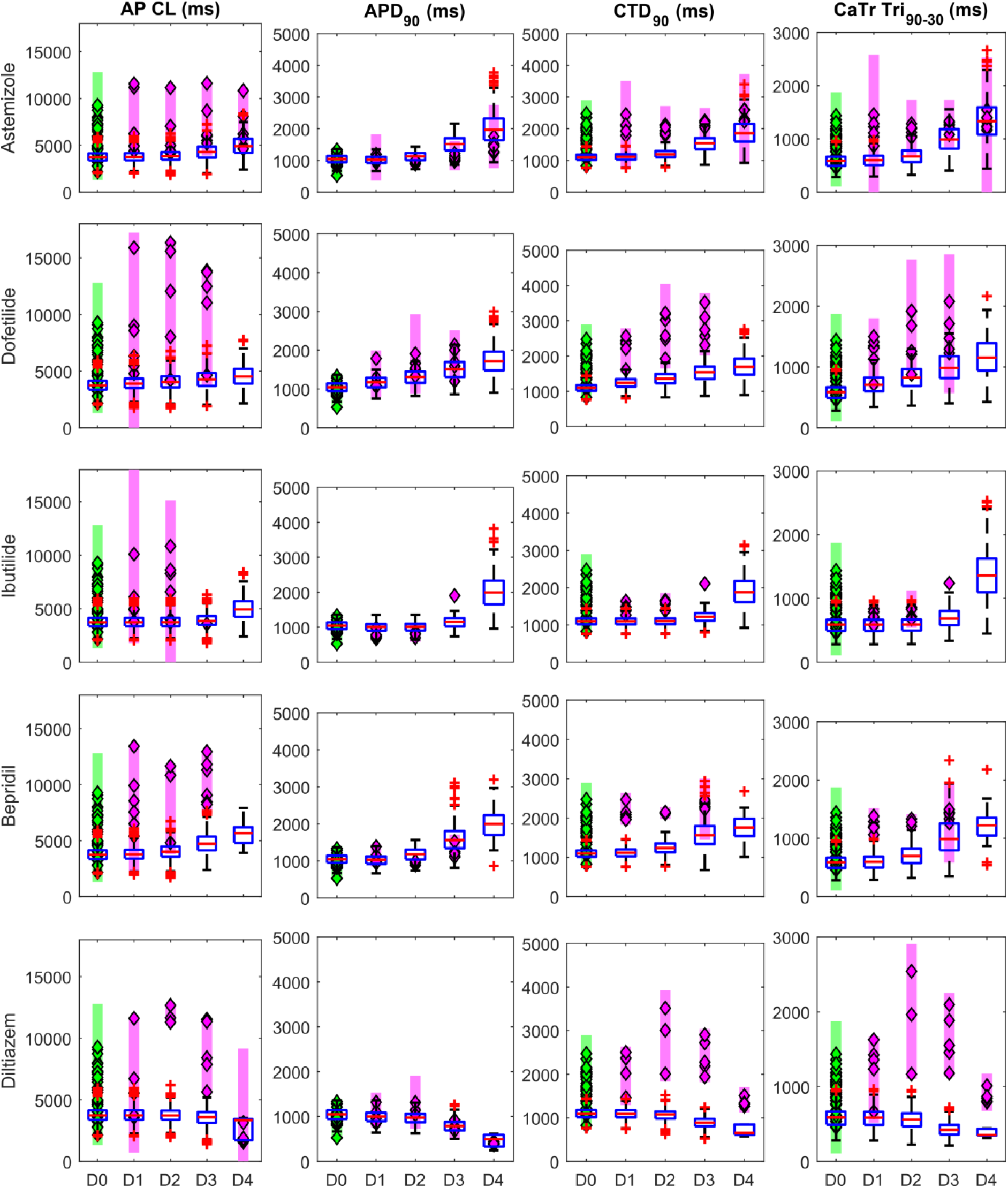
Summary of the drug-induced changes on 4 non-paced AP and CaTr biomarkers in the *in silico* population of hiPS-CMs vs *in vitro* optical recordings. Each line shows results for a different drug, tested at 4 concentrations (D1-D4) and compared to control conditions (D0). Each column corresponds to a different biomarker. In each panel: blue boxplots, simulated biomarkers (Boxplot description as in Figure 3); green diamonds, *in vitro* control biomarkers; purple diamonds, *in vitro* biomarkers following drug applications; green/purple bars, experimental ranges of the *in vitro* data. If no *in vitro* biomarkers are reported for a specific dose, it means that it was not possible computing the biomarkers on the AP and CaTr.

Our *in silico* population, calibrated with optically-recorded biomarkers in control conditions only, reproduces successfully the drug-induced changes in the AP and CaTr biomarkers. If no *in vitro* biomarkers are reported for a specific dose, it means that the drug stopped the spontaneous activity in *in vitro* hiPS-CMs.

The four drugs (astemizole, dofetilide, ibutilide, bepridil), causing a strong I_Kr_ block, induced AP and CaTr prolongation. In particular, simulated APDs, CaTr tRise_0,peak_, and AP and CaTr Tri_90-30_ are well within the experimental ranges. Conversely, simulated AP and CaTr CL and CTDs tend to underestimate the prolongation observed *in vitro*. For diltiazem, a I_CaL_ blocker, simulations reproduced a dose-dependent APD_90_ shortening. However, the CTD_90_ prolongation observed *in vitro* for intermediate doses (D2 and D3) was not captured *in silico*. Table 3 reports the occurrences of drug-induced repolarisation abnormalities and quiescent phenotypes, both in simulations and experiments.

**Table 3.** Drug-induced abnormalities observed in *in silico* vs *in vitro* non-paced hiPS-CMs.

| Drug | Dose | *In silico* |  |  |  |  | *In vitro* |  |  |  |  |  |
| --- | --- | --- | --- | --- | --- | --- | --- | --- | --- | --- | --- | --- |
|  |  | OK | Q | RA | IRR | RESAC | OK | Q | RA | IRR | RESAC | Tachy |
| Astemizole | D1 | 476 | 1 | - | - | - | 6 | - | - | - | - | - |
|  | D2 | 475 | 2 | - | - | - | 6 | - | - | - | - | - |
|  | D3 | 466 | 2 | 4 | 5 | - | 6 | - | - | - | - | - |
|  | D4 | 432 | 2 | 38 | 5 | - | 6 | - | - | - | - | - |
| Bepridil | D1 | 472 | 5 | - | - | - | 5 | - | - | 1 | - | - |
|  | D2 | 466 | 11 | - | - | - | 6 | - | - | - | - | - |
|  | D3 | 365 | 107 | 3 | 2 | - | 6 | - | - | - | - | - |
|  | D4 | 27 | 444 | 6 | - | - | - | 6 | - | - | - | - |
| Diltiazem | D1 | 477 | - | - | - | - | 5 | - | - | 1 | - | - |
|  | D2 | 452 | 25 | - | - | - | 4 | - | 1 | 1 | - | - |
|  | D3 | 204 | 269 | - | 4 | - | 6 | - | - | - | - | - |
|  | D4 | 12 | 444 | 1 | - | 20 | - | - | - | 1* | - | 6* |
| Dofetilide | D1 | 474 | 2 | - | 1 | - | 4 | - | - | 2 | - | - |
|  | D2 | 470 | 2 | - | 5 | - | 5 | - | - | 1 | - | - |
|  | D3 | 466 | 2 | 3 | 6 | - | 5 | - | - | 1 | - | - |
|  | D4 | 461 | 3 | 4 | 9 | - | - | - | 6* | - | - | 2* |
|  | D5 | 455 | 3 | 12 | 7 | - |  |  |  |  |  |  |
|  | D6 | 435 | 1 | 39 | 2 | - |  |  |  |  |  |  |
|  | D7 | 414 | 1 | 59 | 3 | - |  |  |  |  |  |  |
| Ibutilide | D1 | 477 | - | - | - | - | 5 | - | - | 1 | - | - |
|  | D2 | 477 | - | - | - | - | 3 | - | - | 3 | - | - |
|  | D3 | 474 | 2 | - | 1 | - | - | - | 6 | - | - | - |
|  | D4 | 427 | 1 | 47 | 2 | - | - | - | 5 | - | - | 1 |

OK: spontaneous beating with no abnormalities; Q: quiescence; RA: repolarization abnormalities (early afterdepolarizations and/or repolarization failure); IRR: irregular rhythm; RESAC: residual activity; Tachy: tachyarrhythmic oscillations; '-': phenotype not observed; \*: *in vitro* observations showed more than one abnormal phenotype. For dofetilide, D5=10xEFTPC<sub>max</sub>, D6=30xEFTPC<sub>max</sub> and D7=100xEFTPC<sub>max</sub> were tested only *in silico*, to assess if doses higher than D4 would have triggered more abnormalities.

The *in vitro* dataset showed overall less abnormalities in hiPS-CMs in response to drugs than the simulations. A likely reason for this could be that *in silico* results assume single-cell behavior with a wide range of ionic profiles, while syncytial structures were used *in vitro*, where good cell-cell coupling usually has damping effects on pro-arrhythmic behavior. For the drugs inducing AP prolongation (astemizole, dofetilide, ibutilide and bepridil), the abnormalities recorded *in vitro* were single or multiple early-afterdepolarizations (EADs), corresponding to the types A, B and C reported in (13). We also observed 3 cases of tachyarrhythmia (rate of spontaneous oscillations>2Hz), 2 for Dofetilide (D3 and D4, following EADs) and 1 for ibutilide (D4). Finally, 9 cases of irregular rhythm were observed: 4 for Dofetilide (D1, D2 and D3), 4 for ibutilide (D1 and D2) and 1 for Bepridil (D1). For diltiazem, the abnormalities observed *in vitro* were an irregular rhythm at D1, a multiple EAD and irregular rhythm at D2 and a tachiarrhythmic time course at D4 (in 6 out of 6 observations, 1 also with irregular rhythm).

In the simulations for 4 out of 5 tested compounds (astemizole, dofetilide, ibutilide and bepridil) we observed a variety of drug-induced phenotypes, as seen *in vitro* both in our experiments and in (13). Exemplary *in silico* traces are shown in Figure 7 and compared to *in vitro* experiments: single and multiple EADs (panels A, B, C, D), single EADs (panel E, F), repolarization failures (panels G, H) and irregular rhythms (panels I, J, K, L, M, N). Expanded and additional traces are reported in Figure S12 in the Supporting Material. In addition to AP shortening, for diltiazem we observed a residual electrical activity, characterized by low-amplitude oscillations and an EAD (Figure S13 in the Supporting Material).

**Figure 7.**
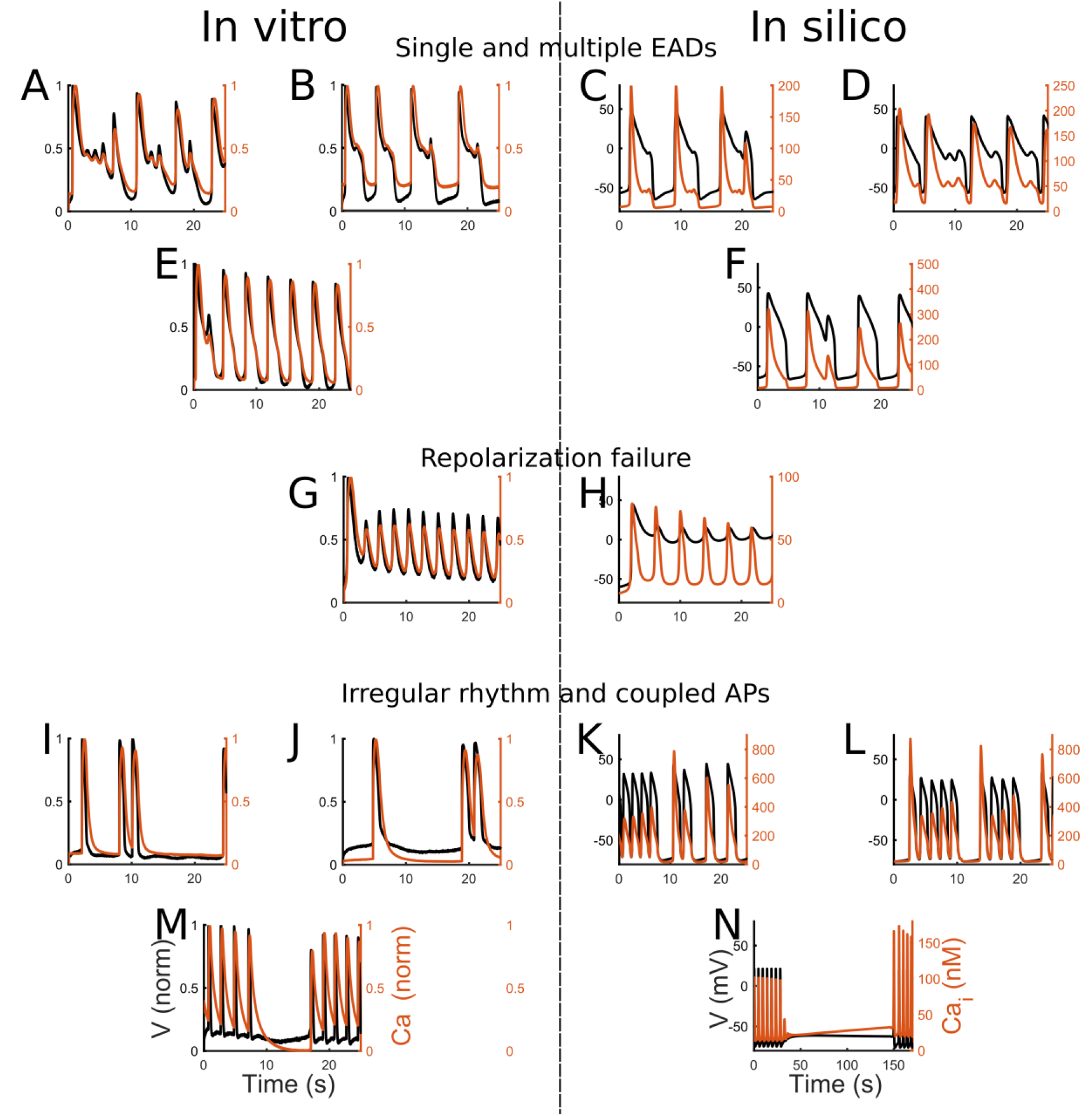
Illustrative abnormalities observed at room temperature during the drug trials *in vitro* (left column) and *in silico* on the population from Figure 5. AP (black) and CaTr (orange). (A, B, C, D) Single and multiple EADs. (E, F) Single EADs. (G, H) Repolarization failure. (I, J, K, L) Irregular rhythms and coupled APs. (M, N) Irregular rhythm, temporary cessation of the spontaneous activity.

For astemizole, *in silico* results reveal 9 abnormalities at D3, and 43 at D4 (mainly EADs and repolarization failures, but also 5 irregular rhythms per dose); the *in vitro* data show dose-dependent increase in pro-arrhythmic markers but no arrhythmia events per se at the tested doses. Again, the syncytial nature of the experimental samples and/or lower temperature may have dampened the arrhythmia events. Nevertheless, the simulation results are in agreement with the fact that at clinical doses, this drug is considered as intermediate risk in (40) on hiPS-CMs and at high risk in the *in silico* drug trials performed in (3) and in CredibleMeds (41). This highlights the value of *in silico* investigations with broader population of models to complement *in vitro* experiments, and ability to cover a wide range of ionic profiles.

Simulations of ibutilide and dofetilide closely agree with the experiments. A dose-dependent increase in abnormalities was seen, typical of drugs classified as high risk in CredibleMeds (41) and in hiPS-CMs in (40). The abnormalities *in silico* are mainly EAD and repolarization failures at the higher doses, together with few cases of irregular rhythm (ibutilide: 1 at D3 and 2 at D4; dofetilide: 1 at D1, 5 at D2, 6 at D3). For dofetilide, at D4 we observed *in silico* only 5 repolarization abnormalities and 9 irregular rhythms, while all 6 *in vitro* recordings showed single or multiple early EADs. Therefore, we tested *in silico* 3 additional doses higher than D4, as in (3), that triggered a considerable amount of EADs (up to 59 EADs/repolarization failures at D7).

Bepridil simulations are in agreement with our *in vitro* experiments. Bepridil’s main effect on hiPS-CMs is the suppression of spontaneous activity in a high percentage of the population (107/477 and 444/477 models, at D3 and D4, respectively). This is consistent with our *in vitro* experiments (6/6 observations at D4 did not produce AP) and with other reports (40). Conversely, only few abnormalities were observed in hiPS-CMs: *in vitro* only 1 irregular rhythm at D1 and *in silico* 5 and 6 abnormalities (2 irregular rhythms and the rest EADs) for D3 and D4, respectively, in agreement also with (40). However, this is in contrast with the high bepridil toxicity observed for adult cells *in vitro* and *in silico*, where it triggers many repolarization abnormalities (3, 41) and might be due to the different expression of ion currents in adult and hiPS-CMs, especially I_CaL_ (13). Therefore, for bepridil only, we tested also the effect of modulating its I_CaL_ blocking power, not changing the drug effect on I_Na_, I_Kr_ and I_NaL_. Figure 8 shows four different models that developed abnormalities with astemizole D4, but not with Bepridil D4 (black traces). However, reducing to half bepridil I_CaL_ blocking power was already enough to trigger EADs. The same behavior was observed by fully inhibiting bepridil I_CaL_ blocking effect.

**Figure 8.**
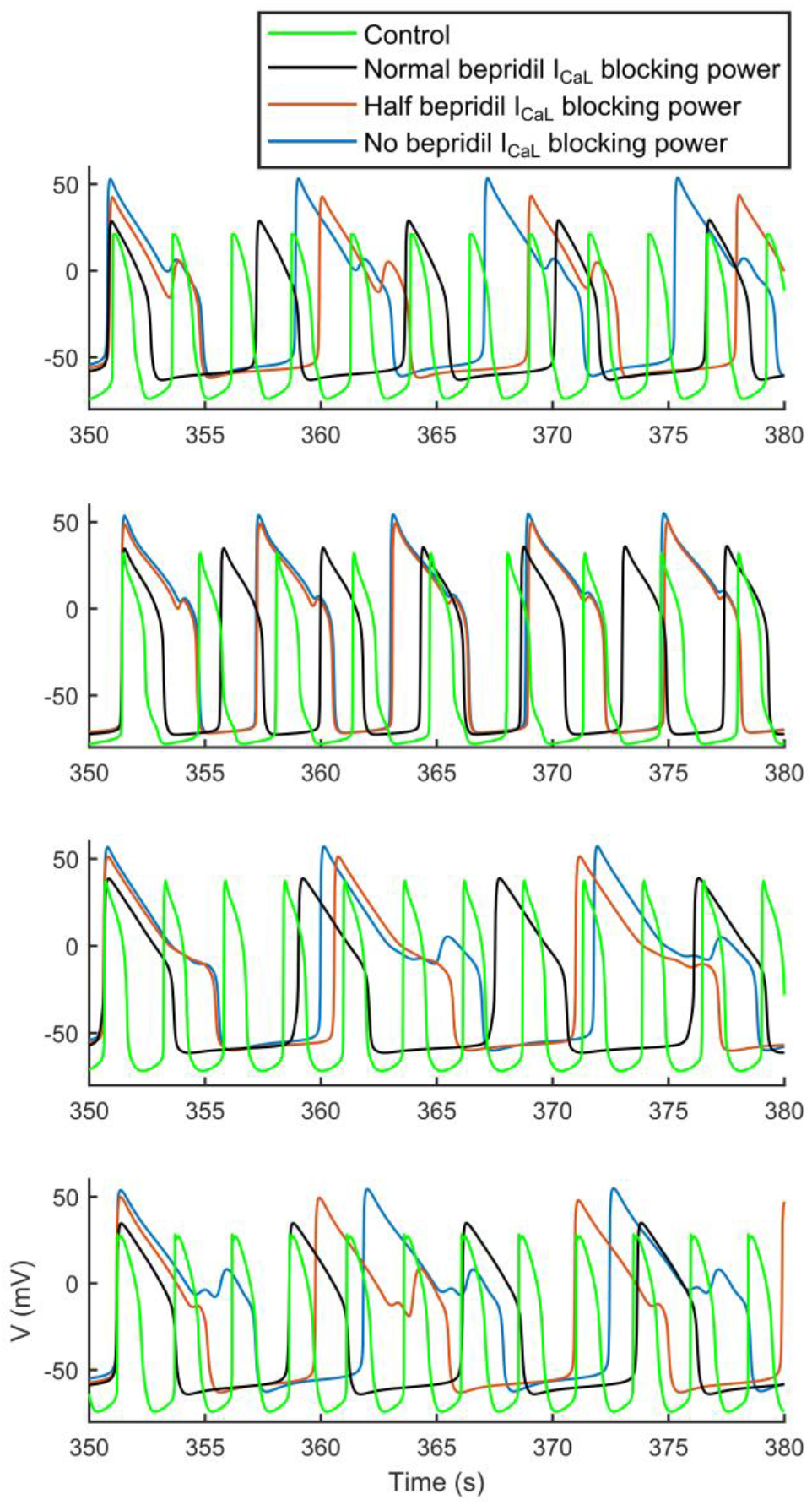
Effect of different I_CaL_ block level during administration of D4 Bepridil. For each of the 4 models (whose control AP are reported in green), we reduced bepridil blocking action of I_CaL_: normal I_CaL_ block (68% block in black); 36% I_CaL_ block (in orange); no I_CaL_ block (in blue). Bepridil effect on the other ion currents was not changed. Drug trials were performed at room temperature.

For diltiazem, we observed *in silico* only 1 EAD at D4 (Figure S13, Panels C), but no tachyarrythmic rhythm, as in our *in vitro* experiments. In fact, most of our models (Table 3) stopped their spontaneous AP, in agreement with what was observed in (40). However, 20 models at D4 showed a strong decrease in AP amplitude (in few cases peaks were recorded below 0mV) and slight increase of frequency (Figure S13, Panels A and B). This low-amplitude oscillations (or residual activity) of the membrane potential were observed in Zeng et al. (42). Zeng et al. demonstrated that such residual electrical activity is due to a residual availability of I_Na_, not fully blocked by drugs specifically designed to mainly block L-type Ca^2+^ channels. It is possible that such abnormal re-activation of I_Na_ may have triggered re-entrant (tachycardic) responses in our multicellular experiments. *In silico* results provide further insights that this residual spontaneous electrical activity is due to a combination of residual I_Na_ (partly blocked by diltiazem, but still able to trigger an AP), strong I_f_ and weak I_K1_ (Table S4 in the Supporting Material, column RESAC).

Simulation studies were used to better understand biophysical mechanisms underlying the drug-induced phenotypes. We observed that the abnormalities induced by astemizole, dofetilide and ibutilide are mainly repolarization abnormalities, while bepridil and diltiazem mainly stopped the spontaneous activity. Table S4 summarizes the ionic parameter differences, the amount of repolarization abnormalities and residual activity at the maximal dose tested *in silico* (D4, except D7 for dofetilide). For the cessation of the spontaneous activity, D3 had more balanced groups for bepridil and diltiazem. We focused our analysis only on those groups containing at least 20 models showing non-sinus rhythm. Models developing EADs and repolarization failures in response to astemizole, dofetilide and ibutilide show weak I_Ks_ and I_K1_ compared to the models not developing such abnormalities, highlighting a reduced repolarization reserve. Also I_pCa_, an outward flow of Ca^2+^ ions is very small, contributing to accumulation of positive charges in the cytosol. Conversely, a different pattern emerged for the models that terminated their spontaneous activity in response to bepridil and diltiazem. They show, compared to the models still developing AP at D3, a strong I_K1_ that stabilizes the resting potential. Furthermore, especially for bepridil, the stronger I_up_ half saturation constant K_up_ reduces the intake of Ca^2+^ by the SERCA pump and therefore the Ca^2+^ available to be released from the sarcoplasmic reticulum, impairing the Ca^2+^handling that it is now an important component of automaticity in the Paci2019 model. For diltiazem, we found that I_Na_ was smaller in models where the drug terminated spontaneous activity compared to the group that still showed spontaneous activity.

## Discussion

Here we demonstrate the integration of human *in silico* drug trials and optically-recorded simultaneous AP and CaTr data from hiPS-CMs for prediction and mechanistic investigations of drug action. We report:

- An improved version of the Paci2018 hiPS-CM model (15) was developed and validated. It better reflects the mechanisms underlying AP automaticity.
- The value of comprehensive high-throughput optical measurements of cellular responses, especially combining AP and CaTr, is demonstrated in refining *in silico* populations of models.
- The predictive power of the experimentally-calibrated population of hiPS-CMs models is demonstrated through *in silico* drug trials on 5 drugs and comparison to *in vitro* datasets.
- Mechanistic insights are gleaned from *in silico* population runs to understand differential responses of hiPS-CM and adult cardiomyocytes to bepridil. Despite observed cardiotoxicity in adult cells (3, 41), *in vitro* experiments, in this dataset as well as in another independent *in vitro* dataset (40), showed low occurrence of proarrhythmic markers in hiPS-CMs. *In silico* trials with the hiPS-CMs models show a wide range of responses to drug action, which complement and explain the *in vitro* experiments.

Research on hiPS-CMs is rapidly developing, with new experimental data becoming available, which in turn serve as a driving force for the constantly evolving computational models to offer more accurate in silico tools to the scientific community. Based on *in vitro* (33) and *in silico* (16) tests, it was identified that our Paci2018 hiPS-CM model (15) did not properly reflect the role of I_NCX_ in automaticity, i.e. no cessation of spontaneous activity was seen in the model as consequence of a strong I_NCX_ block, as suggested by experiments. Therefore, we updated this hiPS-CM model to reproduce the specific mechanisms reported in (16, 33). (Figures 1, S1 and Supporting Material). In addition, the new Paci2019 model also qualitatively simulates the relationship between changes in CL and APD_90_ as consequence of the I_f_ modulation (Figure S16 in the Supporting Material). The model responds to I_f_ augmentation with shorter CL and APD_90_, while I_f_ reduction increases them. In Rast et al. (45), a similar relationship was observed in iCells (CDI) hiPS-CMs field potentials between the inter-beat interval and the field potential duration for ivabradine (for I_f_ reduction) and forskolin (for I_f_ augmentation).

Using the Paci2019 model to construct an *in silico* population based on our *in vitro* optical recordings, we showed that the combination of AP and CaTr biomarkers provides superior calibration, with a better coverage of the biomarker space (Figures 3). It is also interesting that the calibration with AP biomarkers was the most restrictive: AP_CaTr and the AP_only populations contained only 477 and 968 accepted models, respectively, while the CaTr_only population contained over 5,000, many of which were inadequate, e.g. presented extremely short or long APs (Figure S14 in the Supporting Material). Therefore, model calibration exclusively based on CaTr can easily lead to inclusion of more unrealistic models for hiPS-CM. For this cell type, AP biomarkers are preferred to obtain physiological (or semi-physiological) models, while combining both biomarkers clearly refines the calibration. These tests highlight the importance of the calibration process and one of the main advantages of comprehensive records, such as the ones obtained through all-optical cardiac electrophysiology systems like OptoDyCE, that allow the acquisition of AP and CaTr in large populations of cells in their multicellular context.

Figures 6 and S7-11 compare simulated and experimental biomarkers. Of note, the experimental drug trials were not used to calibrate the population of models; yet, the experimentally observed biomarker trends over increasing drug doses, in particular APDs, CTDs and Tri_90-30_, were successfully reproduced. Moreover, for CaTr tRise_0,peak_ and AP and CaTr Tri_90-30_, simulations showed good reproduction of the experimental variability intervals. CTDs were generally underestimated at the various drug doses. A possible reason for this is the fact that in the control population (Figure 3) CTDs are included in the variability ranges, but they cannot cover the higher values. Physiologically-correct *in silico* drug-induced CaTr prolongation (except for diltiazem) was seen, as proven by the overlapping of the *in silico* and *in vitro* CaTr Tri_90-30_. However, the CTD_90_ and CTD_30_ absolute values after drug administration were overall smaller *in silico* than *in vitro*.

We were able to obtain the same abnormality classes (Figure 7) observed in our *in vitro* data and in (13), i.e. single and multiple EADs (panels A, B, C, D, E, F), with the addition of repolarization failure (panels G and H), and irregular rhythms (panels I, J, K, L, M, N). Conversely, the *in silico* models did not show tachyarrhythmias observed e.g. in (13) or in 6 cases in our *in vitro* experiments in response to the highest dose of diltiazem. As discussed previously, these tachyarrhythmias may be syncytium-level events *in vitro* that could not have been captured in the simulations. Furthermore, a common response of the *in silico* hiPS-CMs, especially to administration of diltiazem and bepridil, is the suppression of spontaneous activity. Indeed, diltiazem administration at D3 and D4 also stopped the spontaneous AP in a big portion of our *in silico* population, 269 and 444 models out of 477, respectively. This is in agreement with the *in vitro* diltiazem experiments of 7 out of 15 laboratories involved in the multisite study reported in (40), where 100% of the hiPS-CMs tested did not produce spontaneous AP after administration of 10μM diltiazem (equals to our D4). Furthermore, in 5 laboratories a variable amount (20% - 70%) hiPS-CMs stopped beating. The same effect was observed for bepridil. In fact, as consequence of D3 and D4 bepridil administration, 107 and 444 models out of 477 stopped. Again, this is in agreement with our *in vitro* experiments (no spontaneous AP at D4), and with the experiments of (40) (50% hiPS-CMs stopped spontaneous AP in 4 laboratories (out of 15) with D3 bepridil, and over 80-90% hiPS-CMs in 15 laboratories with D4 bepridil).

It is interesting to note that in our *in vitro* experiments, despite the reliable AP and CaTr duration and triangulation increase, astemizole did not induce abnormalities, while they were observable in 9 *in silico* hiPS-CMs at D3 and 43 at D4. Astemizole is considered an intermediate risk drug in (40) and a high-risk drug both *in vitro* (46) and in the *in silico* drug trials performed in (3). Especially in Blinova et al. (40), 11/15 laboratories observed single and multiple EADs in 100% of their cells at 37°C, in response to 0.1μM astemizole (equivalent to our D4). The absence of EADs in our *in vitro* data (while showing pro-arrhythmic markers such as APD prolongation and increased APD triangulation), may be due to a number of reasons. One possibility is the lower temperature, though temperature-corrected *in silico* hiPS-CMs revealed repolarization abnormalities. Another reason could be potentially higher I_K1_ (and or I_Ks_) in our high-density syncytial preparations compared to other studies.

Overall, hiPS-CMs proved to be an effective *in vitro* and *in silico* model to test drug-induced adverse cardiac effects. Unexpected results *in vitro* and *in silico* for bepridil, considered a highly cardiotoxic drug (3, 41), prompted further investigation. As reported in Table 3, bepridil triggered a very small amount of abnormalities in our *in silico* population. This is in agreement with our *in vitro* experiments and with the tests performed by Blinova et al. (40): in this multisite study, used here for comparison only, bepridil stopped the spontaneous AP in 80-90% hiPS-CMs in all the 15 laboratories at the highest bepridil dose 10μM (in agreement with our simulations); abnormalities were seen only in 2 out of 15 laboratories. Potential reason for this discrepancy can be the higher expression of L-type Ca^2+^ channels observed *in vitro* in hiPS-CMs than in adult cells (13). Blinova et al. (40) state: “*Bepridil is a potent hERG blocker that also blocks L-type calcium and peak and late sodium currents at higher concentrations. High expression levels of calcium ion channels in hiPSC-CMs as compared to primary ventricular tissue may have contributed to more attenuated cellular proarrhythmic effects of the drug as compared to other drugs in the high TdP risk category.*”. We were able to test this idea *in silico*: Figure S15A in the Supporting Material shows I_CaL_ in the original O’Hara-Rudy model of human adult ventricular cell (34) (black trace) and in our hiPS-CM *in silico* population translated to 37°C (cyan traces). We tested if high levels of I_CaL_ could have had a pseudo-protective effect against bepridil in hiPS-CMs, partially compensating the I_Kr_ block, resulting in a milder effect than in cells expressing less I_CaL_ (e.g. adult cardiomyocytes). At room temperature, we tested bepridil D4 on 4 *in silico* hiPS-CMs that showed abnormalities with astemizole, by reducing bepridil I_CaL_ blocking power first to half of its original value and then completely. This resulted in abnormalities in all 4 models (Figure 8), as expected. In addition, these 4 models in control conditions and 37°C showed I_CaL_ higher than the adult one (Figure S15B). Table S2 shows the IC_50_ used for our *in silico* drug trials, taken from (3). Bepridil has the closest I_Kr_ and I_CaL_ IC_50_ among APD-prolonging drugs. Therefore, an I_CaL_ block comparable to I_Kr_ block in condition of highly expressed I_CaL_ could indeed compensate APD prolongation and mask the occurrence of abnormalities, which may have occurred in adult cardiomyocytes (as reported *in silico* in (3, 47)). Our *in vitro* and *in silico* tests show the undeniable value of hiPS-CMs as models for drug testing and how *in silico* simulations could benefit the interpretation of the *in vitro* tests. The hiPS-CMs represent a potentially infinite pool of human cardiomyocytes and can capture key aspects of human cardiac electrophysiology in normal and diseased conditions (genetic mutations). Therefore, they are a great asset to predict the occurrence of adverse drug effects, in a unparallel manner that can be patient-specific.

As all experimental models, the hiPS-CMs are not without limitations. For example, they have different ion current expressions than adult cardiomyocytes, potentially affecting I_Na_, I_CaL_, I_Kr_ and I_Ks_ (see Figure 2 in (13)), i.e. currents for which IC_50_ values are commonly computed. It must be noted that extensive experimental datasets from healthy adult human cardiomyocytes are non-existent due to unavailability of such cardiac tissue. Thus, inferences could only be made based on donor heart-derived human cells (34, 47, 48) or well-studied adult cardiomyocytes from other species. Nevertheless, different ion channel expressions can lead to underestimation (as for bepridil) or overestimation of the actual toxicity of a drug. A variety of optimization approaches are being developed to improve the maturity of the hiPS-CMs and bring them closer to an adult phenotype. These include extracellular matrix optimizations, stimulation protocols, mass transport improvements, alignment, substrate and metabolic function optimizations etc. (49). Such advances can impact positively cardiotoxicity testing.

Overall, commercial hiPS-CMs (e.g. CDI) have demonstrated their utility and superiority to animal models, even in their current state of maturity. Here we show the suitability of optically-recorded data from hiPS-CMs to produce information that empowers *in silico* modelling. With suitably-high acquisition rates, optical data can provide accurate temporal biomarkers for *in silico* models. All available Ca^2+^ data is indeed obtained by optical means; with the development of new small-molecule and genetically-encoded voltage dyes, AP records may completely replace electrical measurements due to their contact-less nature, easy parallelization and ability to measure cell properties in multicellular context. However, absolute values remain a challenge for optical measurements as voltage and Ca^2+^-sensitive dyes are rarely calibrated, i.e. they cannot provide reliable amplitude information for AP or CaTr, i.e. mV or mM. Such absolute values were essential in (19) to calibrate our first hiPS-CM population; in fact, AP peak <57.7mV (19) was included as a biomarker here to avoid unrealistic membrane potentials.

During our *in silico* tests, three limitations emerged. Firstly, CTD_90_, CTD_50_ and CTD_30_ are underestimated during drug administration (Figure 6, rows 1-4). The reason is that the 477 models in the population show relatively short control CaTr despite correct inclusion in the variability ranges by calibration. While the *in silico* CaTr correctly captured the drug-induced trends, they underestimated the changes observed experimentally. The *in silico* CaTr Tri_90-30_ matched well the experimental values, i.e. CaTr triangulation during drug administration was captured. In case of diltiazem (Figure 6, last row) we observed a peculiar behavior of the *in vitro* measurements following drug administration, since the CaTr showed larger CTDs at D2 and D3 than at D1, in spite CTD_90_ shortening for increasing diltiazem doses is clear from D2 to D4. The second limitation is that up to D4 *in silico* dofetilide generated few abnormalities, while D4 dofetilide triggered *in vitro* EADs in all the measurements. We observed already in (22) that to induce a remarkable amount of EADs or repolarization failures in an *in silico* hiPS-CM population, we needed I_Kr_ block>90%. Conversely, D4 dofetilide blocks only 80% I_Kr_. With higher doses, tested in (3), we obtained a considerable increase in AP abnormalities. Finally, we did not observe in our simulations tachyarrhythmias as seen *in vitro* in a few samples, perhaps due to difference in single vs. multicellular behavior. We observed higher spontaneous AP rates, e.g. in irregular rhythms (e.g. in Figure S12, panel I, AP rate goes to 0.59Hz or a cycle of 1700ms) or residual activity in case of diltiazem (in Figure S13, Panel B, rate up to 0.83Hz, corresponding to AP CL of 1200ms). However, we did not observe AP rates greater than 2Hz.

## Conclusions

In conclusion, this work supports the use of high-content, high-quality all-optical electrophysiology data to develop, calibrate and validate computer models of hiPS-CM for *in silico* drug trials. We report that simultaneously-acquired AP and CaTr enhance the model calibration process to obtain a final population that better reflects the experimental recordings. Our population was able to reproduce the effect of 5 different compounds, including the drug-induced abnormalities observed *in vitro*. *In silico* models constrained by *in vitro* data can be used to expand the parameter space of the investigations and to glean mechanistic insights into drug action. Finally, our simulations highlight the importance of being aware and taking into account potential differences in ionic currents between hiPS-CMs and adult cardiomyocytes, which could result in differences between *in vitro/in silico* hiPS-CMs and *in vivo* outcomes for specific compounds.

## Author Contributions

AK and EE recorded and analyzed the optical *in vitro* data. MP and SS designed the Paci2019 hiPS-CM model. MP, EP, SS, JH, BR and EE designed the *in silico* tests on the populations of models. MP implemented the models and software tools used to produce and analyze the *in silico* data. MP, EP and SS analyzed the *in silico* data. All authors contributed to the writing and reviewed the manuscript.

## Acknowledgements

The authors thank Dr. Jussi Koivumäki for sharing his code and for the fruitful discussion about the updates to the hiPS-CM model. The authors wish also to acknowledge CSC – IT Center for Science, Finland, for generous computational resources. MP was supported by the Academy of Finland (decision number 307967). EP and BR were supported by an NC3Rs Infrastructure for Impart Award (NC/P001076/1), a Wellcome Trust Senior Research Fellowship in Basic Biomedical Sciences (100246/Z/12/Z, 214290/Z/18/Z), EPSRC Impact Acceleration Awards (EP/K503769/1), the CompBioMed project (European Commission grant agreement No 675451), the Oxford BHF Centre of Research Excellence (RE/08/004/23915, RE/13/1/30181) and the TransQST project (Innovative Medicines Initiative 2 Joint Undertaking under grant agreement No 116030, receiving support from the European Union’s Horizon 2020 research and innovation programme and EFPIA). EE was supported by the National Institutes of Health (R01HL144157) and the National Science Foundation (1827535 and 1830941).

## Supporting Citations

References (50–55) appear in the Supporting Material.

